# Bone mineral density affects tumor growth by shaping microenvironmental heterogeneity

**DOI:** 10.1101/2024.07.19.604333

**Authors:** Matthew A. Whitman, Madhav Mantri, Emmanuel Spanos, Lara A. Estroff, Iwijn De Vlaminck, Claudia Fischbach

## Abstract

Breast cancer bone metastasis is the leading cause of mortality in patients with advanced breast cancer. Although decreased mineral density is a known risk factor for bone metastasis, the underlying mechanisms remain poorly understood because studying the isolated effect of bone mineral density on tumor heterogeneity is challenging with conventional approaches. Here, we investigate how bone mineral content affects tumor growth and microenvironmental complexity *in vivo* by combining single-cell RNA-sequencing with mineral-containing or mineral-free decellularized bone matrices. We discover that the absence of bone mineral significantly influences fibroblast and immune cell heterogeneity, promoting phenotypes that increase tumor growth and alter the response to injury or disease. Importantly, we observe that the stromal response to matrix mineral content depends on host immunocompetence and the murine tumor model used. Collectively, our findings suggest that bone mineral density affects tumor growth by altering microenvironmental complexity in an organism-dependent manner.

## INTRODUCTION

Bone matrix is crucial for the normal function of the skeletal system and also plays a significant role in the development of pathologies such as osteoporosis and breast cancer bone metastasis^1,2^. Both conditions lead to the weakening of bones through osteolytic degradation, resulting in reduced bone density, chronic pain, increased fragility, and ultimately, worse clinical prognosis^2^. It is widely appreciated that phenotypic changes of osteoblasts, osteocytes, and osteoclasts govern pathological bone loss and thus, bone matrix changes^3^, but it is much less well known how bone matrix changes alter the recruitment and activity of stromal and cancer cells, due in part to a lack of model systems that can separate the effects of changes in bone mineral density from other factors.

Healthy bones are hierarchically structured and primarily composed of hydroxyapatite (HA)-embedded type I collagen matrix. Approximately 70% of the matrix in bone is normally composed of HA, but HA content declines with age, diet, and disease^4–6^; i.e., conditions independently associated with increased risk of bone metastasis^7–9^. Moreover, pathologic fractures result in the formation of hypomineralized, collagen type-I-rich bone matrix^10^, while increasing metastatic colonization around the injury site^11^. As bone resident cells can sense and respond to bone matrix changes and secrete factors that influence the recruitment of other cell types, pathological alterations in HA content are likely to influence cellular phenotypes and composition of the skeletal microenvironment and thus, bone pathologies. Indeed, microenvironmental heterogeneity is well-established to affect tumor initiation, growth, and immunity^12–14^, but it is much less clear how reduced bone matrix mineral content controls this interplay and which effects these changes have on disease progression. Elucidating these connections is crucial for advancing prophylactic strategies to interfere with bone pathologies including metastasis.

While *in vitro* studies using HA-mineralized biomaterials demonstrate that bone-resident cells are responsive to variations in mineral content^15,16^, these systems fail to mimic the complex interplay between various cell types *in vivo*. Vice versa, *in vivo* studies enable analysis of cellular complexity in bone, but cannot conclusively determine how mineral content affects these results. For example, osteomalacic hypophosphatemic (hyp) or vitamin D deficient (VDR) mice have both hypomineralized bones and altered immune cell function^17–19^. Whether or not these observations are functionally linked is unclear due to accompanying systemic effects^20,21^. To better understand the isolated effect of HA on cell behavior, we developed protocols to selectively control mineral content in physiologic bone matrices^2,22^ and deconvolve the effect of bone matrix on cancer cells from the effect of bone resident cells. Using these scaffolds *in vitro*, we have previously found that breast cancer cells alter mechanosignaling and adapt their phenotype in response to varying HA content^2^. Nevertheless, how bone matrix mineral content regulates microenvironmental complexity *in vivo* and which effect these changes have on cancer progression is unknown.

Single-cell RNA-sequencing has been used previously to analyze the holistic cellular phenotypic response to biomaterials^23^, disease^24^, and development^25^. In this study, we performed single-cell RNA-sequencing on implanted decellularized bovine bone scaffolds in which the mineral was either maintained at physiological levels or removed to simulate scenarios of impaired bone mineralization as, for example, present during aging^4^, injury^11^, or metastasis^26^. Using this approach, we explored the heterogeneous stromal response to varied bone mineral content in both an immunocompromised and immunocompetent syngeneic mouse model in the presence and absence of cancer cells. Collectively, our study provides new perspectives on the impact of matrix mineralization on microenvironmental complexity and resulting consequences on tumor growth. We additionally contribute new single-cell atlases of cellular response to bone matrix-derived biomaterials, which are widely pursued to repair bone defects in the clinic. Findings from our work hold implications for future research on bone metastasis and contribute to the growing body of knowledge surrounding the systemic response to biomaterial implants.

## RESULTS

### Bone matrix mineral content slows early tumor growth without broad effects on host cell recruitment

To investigate the effect of bone matrix mineral content on cellular heterogeneity, we produced 6 mm diameter, 1 mm thick scaffolds from neonatal bovine trabecular bone as previously described^2^. Scaffolds were decellularized and then either used as matrices with physiological levels of mineralization (M-Bone) or subjected to ethylenediaminetetraacetic acid (EDTA)-based demineralization to yield demineralized scaffolds (DM-Bone) (**Fig. 1A, Methods**). Characterization of the M-Bone versus DM-Bone scaffolds by compression test and microcomputed tomography confirmed that EDTA demineralization reduced the bulk elastic modulus of scaffolds (**Fig. 1B**) and completely removed inorganic mineral matrix components in DM-Bone scaffolds (**Fig. 1C**), respectively. Scanning electron microscopy (SEM) further validated that bone matrix macro- and microstructure were unaffected by EDTA-based demineralization consistent with our previous work^2^ (**Fig. 1D**). In addition, the fibrillar structure of collagen I, the principal organic component of bone matrix, was comparable in both scaffold systems as polarized-light microscopy of Picrosirius Red-stained samples did not reveal differences between both conditions^27^ (**Supp.** Fig. 1A).

**Figure 1.**
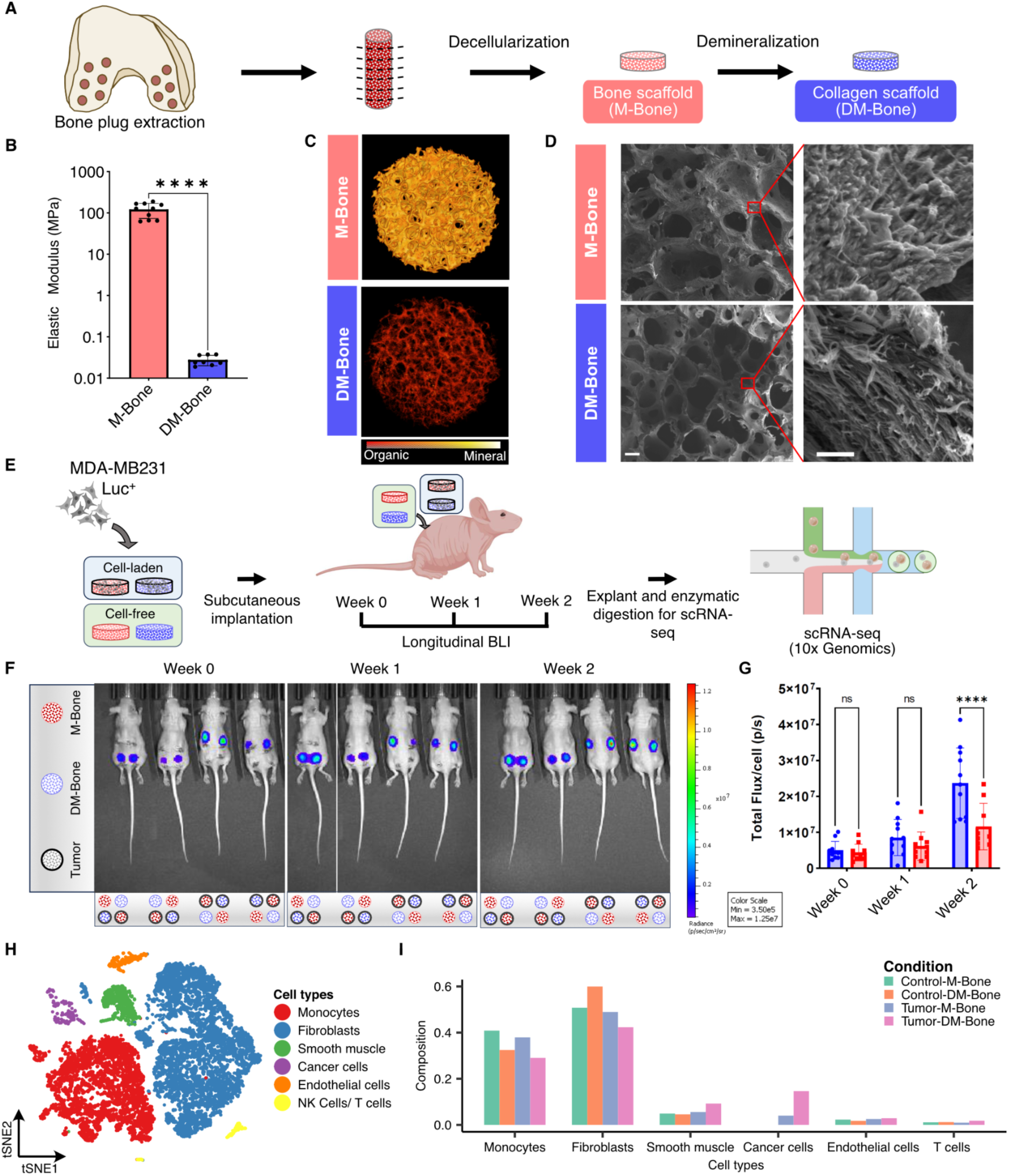
Bone matrix mineral content alters tumor growth without global effects on host cell recruitment. **A)** Schematic showing the process of generating implantable 1-2 mm long, 6 mm diameter bone scaffolds. **B)** Compression testing of M-Bone and DM-Bone scaffolds indicates differences in elastic modulus. (*P=0.0332, **P=0.0021, ***P=0.0002, ****P<0.0001) **C)** Representative microcomputed tomography (micro-CT) images of M-Bone and DM-Bone scaffolds. Pseudocolor indicates mineral content. **D)** Representative SEM images of M-Bone and DM-Bone scaffolds. Scale bar = 200 μm (left) and 2 μm (right). **E)** Schematic showing experimental design for single-cell transcriptomics experiments using luciferase-expressing MDA-MB231 triple negative human breast cancer cells seeded onto M-Bone and DM-Bone scaffolds and implanted into female athymic nude mice. **F)** Longitudinal bioluminescent imaging (BLI) and quantification of luciferase-expressing tumor cells on implanted M-Bone and DM-Bone scaffolds. (****P<0.0001) Pseudocolor indicates radiance pixel intensity between 3.5 x 10^5^ and 1.25 x 10^7^ p s−1 cm−2 sr−1. **G)** Bar plot showing comparison of normalized flux of MDA-MB231 tumor cells on M-Bone and DM-Bone scaffolds at three time points post implantation in athymic nude mice. **H)** t-Distributed Stochastic Neighbor Embedding (t-SNE) map of 16,973 single-cell transcriptomes clustered by gene expression and colored by the labeled cell types. **I)** Bar plot showing relative proportion of various cell types across the four experimental conditions.

To examine the naïve stromal response to bone matrix mineral content, we implanted M-Bone and DM-Bone scaffolds subcutaneously (s.q.) onto the dorsal flanks of 6-8-week-old female immunocompromised athymic nude mice. In these mice, we also implanted M-Bone and DM-Bone scaffolds pre-seeded with luciferase-expressing MDA-MB231 breast cancer cells, a well-accepted and widely used model to study bone metastasis^28,29^ **(Fig. 1E, 1F, Methods)**. As the phenotype of stromal cells recruited during the early-stages of tumor initiation is a critical determinant for the severity of lesion formation^5,30^, we explanted the different scaffolds two weeks after implantation. This protocol enabled assessment of stromal composition following the conclusion of the initial wound healing response^31^ and mimicked early stages of tumor formation. We used longitudinal bioluminescent imaging (BLI) to assess tumor growth weekly. Next, the implants were removed, enzymatically digested, and phenotypically assessed via single-cell RNA-sequencing (scRNA-seq) **(Fig. 1E)**. We observed that immediately after implantation, and up to one week after implantation, bioluminescence did not differ between scaffold types. After two weeks, the luminescence of tumors forming on DM-Bone scaffolds (Tumor-DM-Bone) was significantly higher than on M-Bone scaffolds (Tumor-M-Bone). This observation indicates that reduced bone mineral content promotes tumor growth. **(Fig. 1F, 1G).** After explantation, we generated scRNA-seq data for 16,973 cells from M-Bone and DM-Bone scaffolds implanted with and without tumor cells (**Methods, Fig. 1H, Supp.** Fig. 1A). The single-cell transcriptomes from these different implants identified six distinct cell types for each group, including cancer cells and stromal cells such as endothelial cells, fibroblasts, smooth muscle cells, monocytes, and T cells/NK T cells. As expected, tumor cells were only present in explants of scaffolds pre-seeded with tumor cells. At the study endpoint, more tumor cells were isolated from DM-Bone scaffolds relative to mineral-containing scaffolds, consistent with our BLI data (**Fig 1H, 1I and Supp.** Fig. 1B). Regardless of whether implants contained tumor cells, stromal cells were the most abundant class of cells in the tumor microenvironment. Among stromal cells, fibroblasts and monocytes constituted the two most abundant cell types, with relative proportions that were roughly equal across conditions. Therefore, we investigated these populations in more detail next.

### Bone matrix mineral content regulates the phenotype of recruited fibroblasts

Fibroblasts are intrinsically heterogeneous and critical regulators of both physiological and pathological tissue remodeling. Depending on their phenotype, fibroblasts can direct a profibrotic or wound-healing response to a biomaterial implant with functional consequences for immune cell phenotype and implant engraftment^32,33^. In a tumor, fibroblasts assume a variety of phenotypes, known as cancer-associated fibroblasts (CAFs), that can be tumor-suppressive through the secretion of growth-inhibitory and immunoregulatory cytokines. Conversely, they may also promote tumor outgrowth and invasion through the secretion of growth factors, matrix metalloproteinases (MMPs), and extracellular matrix (ECM) proteins^34–36^. Given this plasticity, and because fibroblasts represented the most abundant host cell type in explants across all conditions, we characterized the fibroblast phenotypes that were associated with the different bone scaffolds in the presence or absence of tumor cells. Unsupervised clustering of single-cell transcriptomes from all four scaffold conditions yielded four distinct subpopulations of fibroblasts that were distributed differently based on mineral content of the scaffold and whether tumor cells were present (**Fig. 2A-2C**). Two distinct clusters, *S100a4*+ and *Saa3*+ fibroblasts, were enriched in M-Bone scaffolds in both tumor and tumor-free conditions. A third cluster of *Col8a1*+ *Mfap4*+ fibroblasts was enriched in tumor-free DM-Bone scaffolds relative to M-Bone scaffolds but almost entirely absent in all tumor-containing explants **(Fig. 2A, 2B and Supp.** Fig. 2A). In contrast, a fourth cluster of *Acta*+ *Tagln*+ fibroblasts was enriched in Tumor-DM-Bone scaffolds relative to Tumor-M-Bone scaffolds, but largely absent in all tumor-free explants.

**Figure 2.**
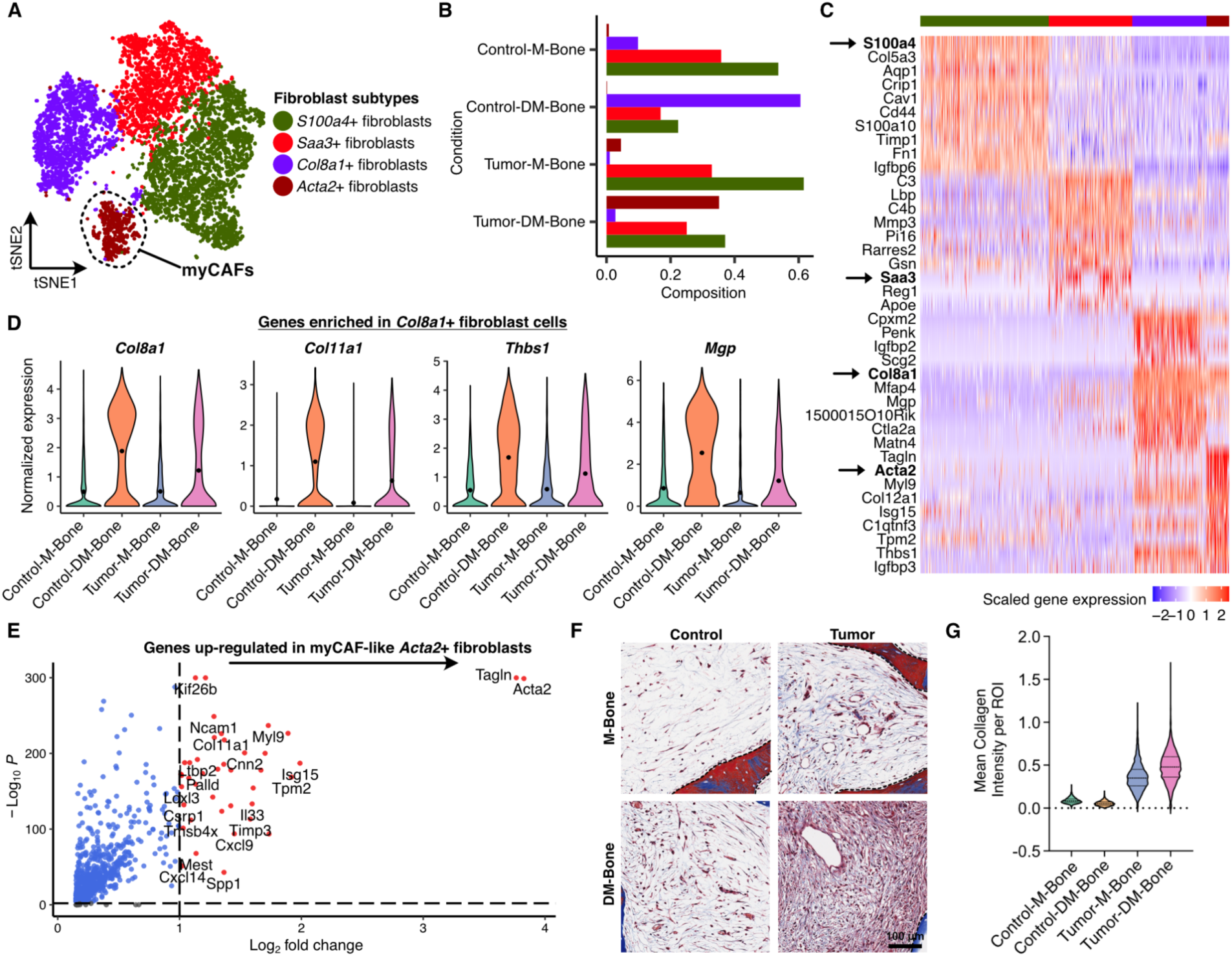
Fibroblast phenotype is altered by bone matrix mineral content and tumor cells. **A)** t-Distributed Stochastic Neighbor Embedding (t-SNE) map of 7,935 fibroblast single-cell transcriptomes clustered by gene expression and colored by the fibroblast clusters. **B)** Bar plot showing relative proportion of various fibroblast cell clusters across the four experimental conditions. Colors represent the fibroblast subtypes as shown in Fig. 2A. **C)** Heat map showing the log-normalized and scaled expression of top-ten differentially expressed genes in each fibroblast cluster. Colors in the color bar on top represent the fibroblast subtypes as shown in Fig. 2A. **D)** Violin plots showing the log-normalized expression of genes upregulated in *Col8a1*+ fibroblast cells enriched on tumor-free demineralized bone scaffolds (Control-DM-Bone). **E)** Volcano plot of differential gene expression analysis showing genes upregulated in myCAF-like *Acta2*+ fibroblasts relative to all other fibroblasts. **F)** Representative Masson’s Trichrome images of explanted tumor-free and tumor-containing M-Bone and DM-Bone scaffolds and **(G)** Analysis of collagen pixel intensity by mean intensity per region of interest. Scale bar = 100 μm, dashed line denotes scaffold.

To isolate the effect of matrix mineral content on fibroblast phenotype in the absence of tumor cells, we analyzed *Col8a1*+ cells as they were much more abundant in all tumor-free explants and highly enriched in DM-Bone scaffolds as compared to M-Bone scaffolds. Interestingly, these cells not only had elevated expression of collagen *Col8a1* but also *Col11a1* as well as matricellular glycoprotein gene thrombospondin, suggesting that decreased bone mineral content induces a matrix-remodeling phenotype in fibroblasts **(Fig. 2D)**. We also observed elevated expression of matrix gla protein (*Mgp)* in these cells, which inhibits pathological calcifications in heart valves and arteries^37^, while also regulating healthy bone formation and activity of bone morphogenetic protein 2 (BMP-2) (**Fig. 2D**)^38,39^. This suggests reduced bone mineral content upregulates factors involved in matrix remodeling, a finding that was supported by differential gene expression and gene ontology analysis of fibroblast cells on tumor-free scaffolds. More specifically, we found that genes associated with bone matrix remodeling including genes associated with bone formation, biomineralization and wound healing were enriched in fibroblasts in tumor-free DM-Bone relative to M-Bone scaffolds (**Supp Fig. 2B-2D**). Last, we analyzed the *Acta2*+ fibroblast cell cluster resembling previously defined cancer-associated fibroblast-Bs (CAF-Bs) and myofibroblastic cancer-associated fibroblasts (myCAFs)^40–42^ (**Fig. 2A, 2C, and 2E**). This cluster was highly enriched in DM-Bone scaffolds as compared to M-Bone scaffolds (**Fig. 2A**) and characterized by high expression levels of the canonical myofibroblast markers alpha-smooth muscle actin (*Acta2*), transgelin (*Tagln*), and myosin light chain 9 (*Myl9*) in addition to the tropomyosin gene *Tpm2*, and matrix-remodeling gene *Mmp13* **(Fig. 2E).**

To validate that mineral affects myCAF phenotype, we performed histology on tumor sections. Increased Masson’s trichrome staining of tumors grown in DM-Bone scaffolds supports that lack of bone mineral induces a collagen-depositing myCAF phenotype **(Fig. 2F, G).** While *Acta2*+ fibroblasts were detected in both tumor-free and tumor-containing explants, considerably more *Acta2*+ cells were detected in tumors growing within DM-Bone scaffolds (Tumor-DM-Bone) relative to all other conditions providing additional evidence for our conclusion **(Supp.** Fig. 2E). Collectively, our data indicate that fibroblast populations are relatively similar in tumor-free and tumor-containing scaffolds as long as mineral is present. In the absence of mineral, however, fibroblasts change dramatically, and these changes are further enhanced by the presence of tumor cells likely shaping the microenvironment in a manner that contributes to increased tumor outgrowth.

Myeloid cells represented the second largest population of cells in the different explants **(Supp.** Fig. 3A). Unsupervised clustering yielded four distinct subpopulations of macrophages that were distributed differently based on mineral content of the scaffold and whether tumor cells were present **(Supp.** Fig. 3A-C). Most macrophages in the M-Bone scaffolds were unactivated *Cd163*+ cells, whose population size decreased in DM-Bone scaffolds regardless of tumor status. Importantly, we also identified two distinct phenotypes of activated monocytes that were enriched on DM-Bone scaffolds relative to M-Bone scaffolds, but whose specific phenotype was further dictated by the presence of tumor cells. Specifically, macrophages in tumor-free conditions expressed tumor necrosis factor (*Tnf*), a master regulator of inflammation and proliferation, and *Ccl2* which is increased by NF-κB signaling and associated with an inflammatory phenotype. These *Tnf*+ macrophages were more abundant in DM-Bone scaffolds relative to M-Bone scaffolds. In contrast, tumor-containing explants were almost devoid of *Tnf*+ macrophages. Macrophages in tumor-containing scaffolds were enriched for *Cd72*+ proinflammatory macrophages in DM-Bone versus M-Bone scaffolds but were almost absent in tumor-free scaffolds (**Supp.** Fig. 3C). We also captured populations of *Cd9*+ macrophages and dendritic cells that did not change significantly across conditions (**Supp.** Fig. 3A-C). Although macrophage phenotypes appeared to be responsive to the matrix, typical markers of tumor-associated macrophages, such as *Nos2*, *Cd274*, *Mmp13*, and *Cxcl3,* were not detected in any of our tumor-containing scaffolds **(Supp.** Fig. 3D).

Although tumor growth was consistently increased in scaffolds devoid of mineral, we observed minimal transcriptional changes in captured tumor cells **(Supp.** Fig. 4A, B). Thus, scaffold-dependent changes to the microenvironment, including altered fibroblast phenotypes, may be more important for tumor growth at this early stage than the phenotype of the tumor cells themselves. This may be particularly important as myofibroblasts resembling those detected in our murine samples are a significant component of human bone metastases (**Supp.** Fig. 5). As the aforementioned experiments were performed with human breast cancer cells in immunocompromised mice to mimic conventionally used mouse models of bone metastasis^43^, it is possible that the absence of a fully functional immune system may have contributed to this observation. Yet, dysregulated bone formation and mineralization are accompanied by changes in immune cell function^17,18^. Thus, we next evaluated how bone mineral content affects immune cell types in immunocompetent mice implanted with syngeneic triple-negative mammary cancer cells.

### Bone matrix mineral content regulates the immune microenvironment

Immune cells play an important role in bone formation and remodeling, and changes of bone materials properties regulate immune cell function^44–46^. For example, immune cells prepare bone niches for subsequent tumor cell colonization^47^, while bone mineral properties, including mineral particle size or crystallinity influence the phenotype of immune cells including dendritic cells^48^ and macrophages^49,50^. As tumor-resident immune cells such as tumor-associated macrophages (TAMs) or tumor-associated neutrophils (TANs) can either promote or suppress tumor growth depending on their specific microenvironmental context^51–53^, it is important to understand which role bone mineral content may play in this process.

To determine how bone matrix mineral content alters stromal cell recruitment in mice with a fully functional immune system, we implanted tumor-free and tumor-containing M-Bone and DM-Bone scaffolds s.q. into the dorsal flanks of 6-8 week old female Balb/C mice **(Fig. 3A).** For the tumor condition M-Bone and DM-Bone scaffolds were seeded with syngeneic luciferase-eGFP expressing triple-negative 4T1.2 mouse metastatic mammary cancer cells prior to implantation. Similar to the studies described above, we tracked tumor cell growth longitudinally and removed the implants after two weeks for single-cell transcriptomics. In comparison to the xenografted human MDA-MB-231 cells, tumor development was much more rapid, BLI signal was unaffected by mineral content of the implants **(Fig. 3B, 3C)**, and tumors were characterized by limited desmoplasia and significant necrosis (**Fig. 3D)**. Consistent with these observations, the parental cell line of the 4T1.2 has been characterized by its rapid growth and granulocytosis^54^, where the dominant microenvironmental phenotype is characterized by necrosis resulting from aggressive tumor growth.

**Figure 3.**
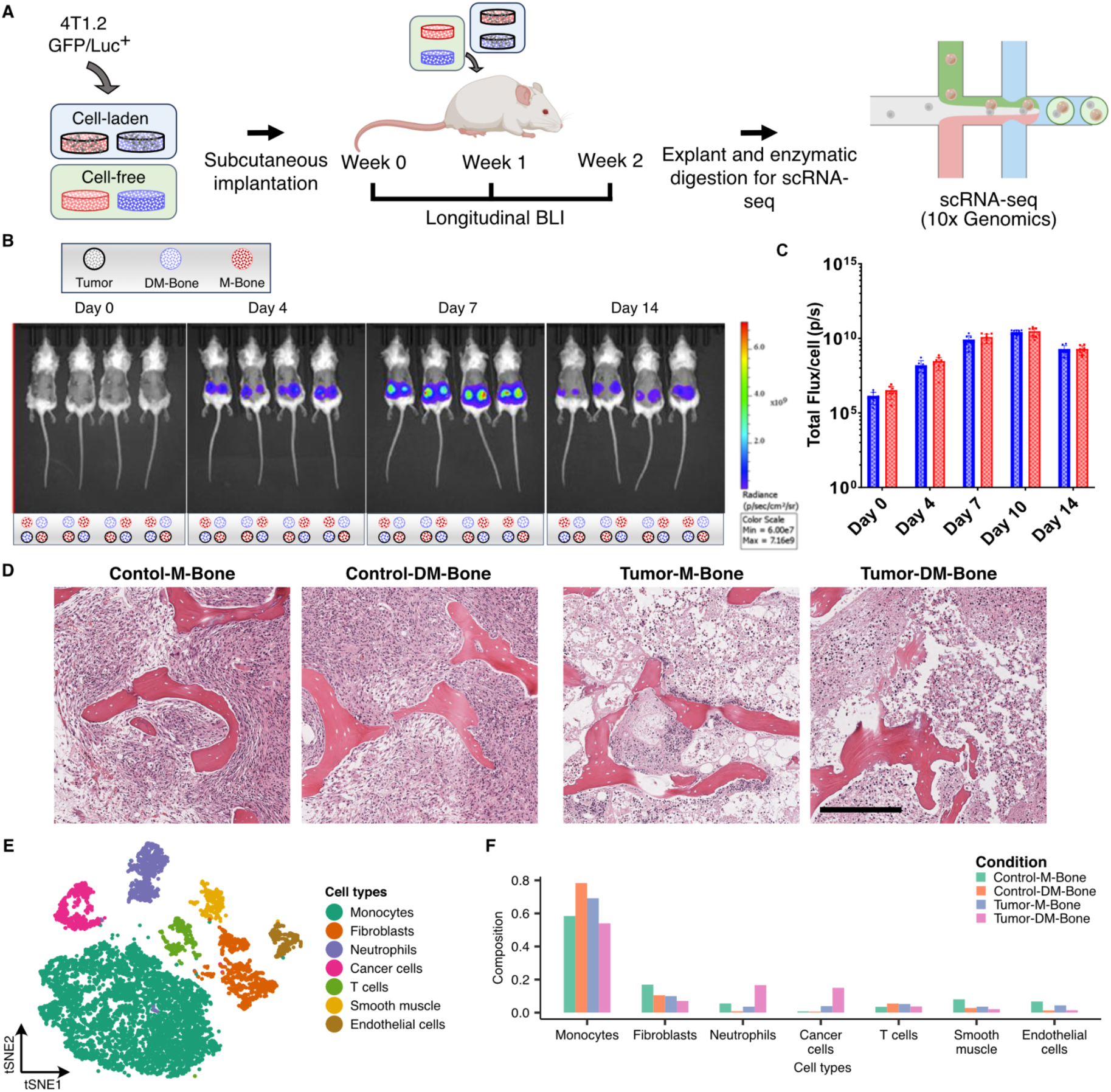
Bone matrix mineral content regulates stromal cell recruitment in immunocompetent mice. **A)** Schematic showing experimental design for single-cell transcriptomics experiments using luciferase-expressing 4T1.2 bone tropic triple negative murine breast cancer cells implanted in Balb/C mice on M-Bone and DM-Bone scaffolds. **B)** Longitudinal bioluminescent imaging (BLI) of luciferase-expressing tumor cells seeded onto M-Bone and DM-Bone scaffolds and implanted s.q. into female Balb/C mice. Pseudocolor indicates radiance pixel intensity between 6.00 x 10^7^ and 7.16 x 10^9^ p s−1 cm−2 sr−1. **C)** Bar plot showing comparison of normalized flux of 4T1.2 tumor cells on M-Bone and DM-Bone scaffolds at three time points post implantation in Balb/C mice. **D)** Representative H&E images of tumor-free and 4T1.2 tumor-containing M-Bone and DM-Bone scaffolds. Scale bar = 300 μm. **E)** t-Distributed Stochastic Neighbor Embedding (t-SNE) map of 9,253 single cell transcriptomes clustered by gene expression and colored by the labeled cell types. **F)** Bar plot showing relative proportion of various cell types across the four experimental conditions.

At study endpoint, we generated scRNA-seq data for 9,253 cells from M-Bone and DM-Bone explants containing or lacking tumor cells **(Fig. 3E and Supp.** Fig. 6A). The single-cell transcriptomes represented seven distinct cell types, including tumor cells and stromal cells such as endothelial cells, fibroblasts, smooth muscle cells, monocytes, neutrophils, T cells/ NK T cells, and B cells. **(Fig. 3F and Supp.** Fig. 6B). Interestingly, the number of fibroblasts in all samples was substantially lower in this model compared to the immunocompromised model and to clinical bone metastasis samples^55^. Fibroblasts accounted for only 7%-17% of the total cells, with slightly different proportions across conditions. These results were consistent with previous findings that tumors from MDA-MB231 are inherently more myofibroblastic and desmoplastic than 4T1 tumors^56^. Fibroblast differences between the two mouse models could be driven by innate differences in tumor cell proliferation as 4T1 tumors in Balb/C mice grow much more quickly than MDA-MB231 in immunocompromised mice^57^ possibly affecting the recruitment of host cells via both altered biochemical and biophysical parameters. Interestingly, myeloid cells were the most abundant cell type representing 53%-78% of the total cells across the four conditions (**Fig. 3F**). In addition, we observed a substantial increase in the number of neutrophils on scaffolds with tumor relative to scaffolds without tumors (**Fig. 3F**). Clustering of neutrophil transcriptomes revealed that this difference was largely mediated by an increase of *Ccl3+ Cxcl3+* activated neutrophils expressing *Icam*+ *Cxcr2*-*Sell*-, which are canonical markers of tumor-associated neutrophils (TANs) . These activated TAN-like cells were enriched on DM-Bone scaffolds relative to M-Bone scaffolds in both the absence and presence of tumor cells (**Supp.** Fig. 7A-E). A small increase in the proportion of activated neutrophils on DM-Bone scaffolds, even in absence of tumor cells, suggests that mineral alone can influence neutrophil activation (**Supp.** Fig. 7A**, 7C**). Further analysis of gene markers for N1 (anti-tumor) and N2 (pro-tumor) type neutrophils suggested that the activated TAN-like cells expressed pro-inflammatory N1-type markers such as *Il1a, Tnf*, and *Ccl3* and low levels of typical N2 markers (**Supp.** Fig. 7F-G). Our results imply that increased tumor growth caused by lack of mineral could enhance the recruitment of activated N1-type TANs, which can promote a tumor-suppressive microenvironment by altering the immune response^52^. As expected, cancer cells were only detected in the tumor samples and represented 3.9% and 15.0% of total cells in M-Bone and DM-Bone explants, respectively. These results are consistent with the immunocompromised system, in which we also detected fewer cancer cells on M-Bone relative to DM-Bone **(Fig 1I, 3E)**. Hence, it is possible that 4T1.2 tumors in Balb/C mice grew more on DM-Bone scaffolds relative to M-Bone, but that these differences were undetectable via BLI due to the significant amount of necrosis.

### Macrophage activation state is altered by bone mineral content

Since monocytes were the most abundant cell type in our dataset and are primarily composed of macrophages, we next analyzed macrophages. Macrophages play important roles in dictating the response to biomaterial implants and are ubiquitous at bone-metastatic sites. Macrophages also represent the most abundant leukocyte population in mammary tumors and play critical and multifaceted roles at each stage of cancer progression^58^. For example, tumor-associated macrophages facilitate neoplastic transformation, tumor immune evasion, and metastasis, but can also drive tumor suppression^58^. To understand how the individual and combined effects of mineral and tumor presence affect the phenotype of macrophages, we reanalyzed the monocytes detected in our tumor-free and 4T1.2 tumor-containing M-Bone and DM-Bone scaffolds **(Fig. 4A, Methods).**

**Figure 4.**
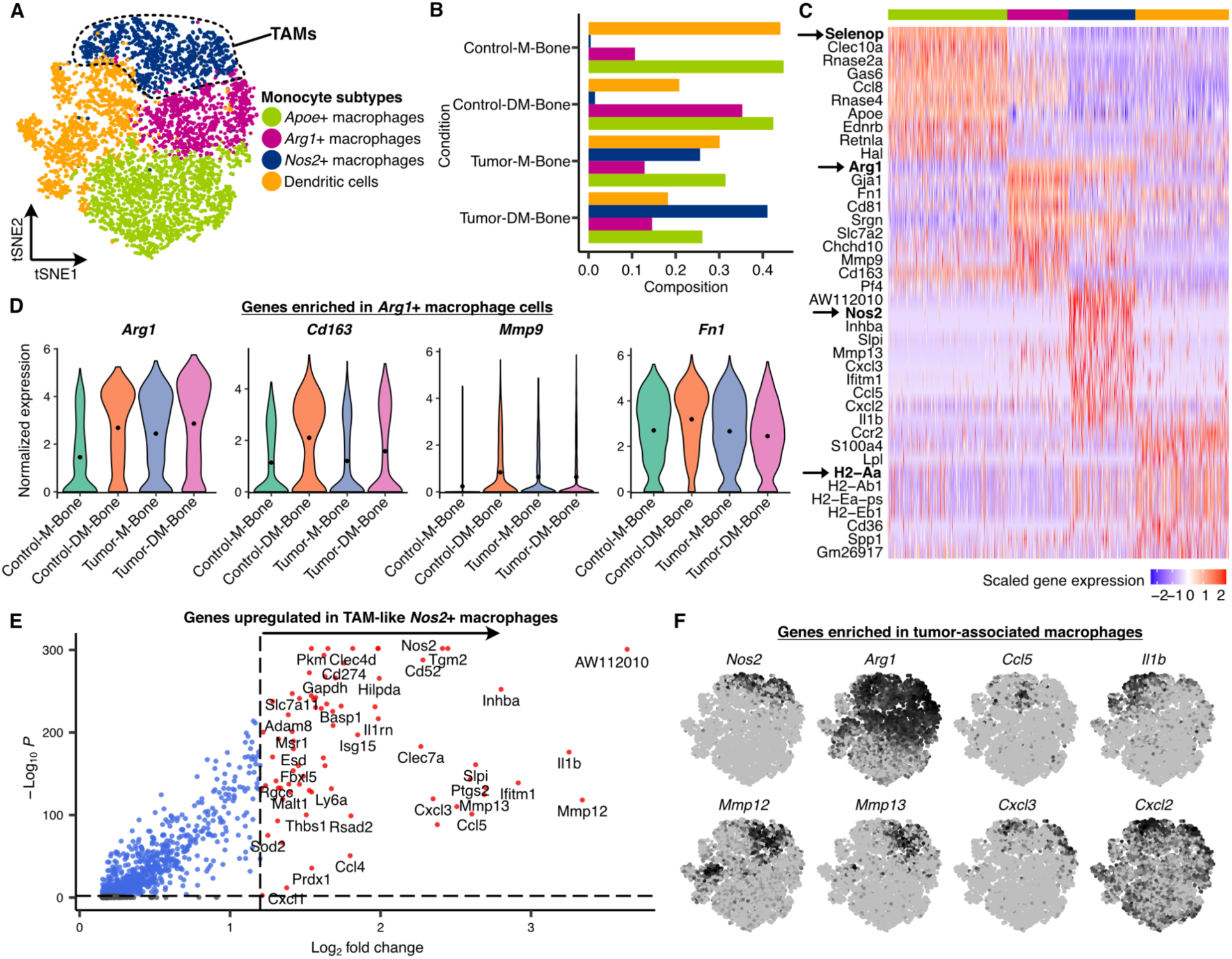
Macrophage activation state is altered by bone mineralization. **A)** t-Distributed Stochastic Neighbor Embedding (t-SNE) map of 5,836 monocyte single-cell transcriptomes clustered by gene expression and colored by the monocyte subtype clusters. **B)** Bar plot showing relative proportion of various monocyte cell clusters across the four experimental conditions. Colors represent the monocyte subtypes as shown in Fig. 4A. **C)** Heat map showing the log-normalized and scaled expression of top-ten differentially expressed genes in each monocyte cluster. Colors in the color bar on top represent the monocyte subtypes as shown in Fig. 4A. **D)** Violin plots showing the log-normalized expression of genes enriched in monocyte cells on DM-Bone (top row) and on M-Bone (bottom row) scaffolds. **E)** Volcano plot of differential gene expression analysis showing genes upregulated in TAM-like *Nos2*+ macrophages relative to all other macrophage cells. **F)** t-SNE feature plots showing expression of genes enriched in tumor-associated macrophages.

Unsupervised clustering of monocytes revealed three distinct macrophage clusters: *i) Selenop*+ *Apoe*+ macrophages were similarly abundant on M-Bone and DM-Bone but were slightly decreased in the presence of a tumor suggesting that this subpopulation is not responsive to mineral content but regulated by tumor cells. *ii) Arg1*+ *Gja1*+ macrophages were highly enriched in tumor-free scaffolds devoid of mineral, but mineral had no effect when tumor cells were present. Hence, this subpopulation could regulate bone remodeling as a function of bone mineral content in tumor-free conditions. Vice versa, *iii) Nos2*+ *Inhba*+ macrophages were detected almost exclusively in tumor-containing scaffolds and were enriched on Tumor-DM-Bone vs. Tumor-M-Bone scaffolds. These results imply that this subpopulation may affect tumor progression differentially depending on bone mineral content **(Fig. 4B)**. We also detected a small population of *H2-Aa*+ *Ccr2*+ dendritic cells that was slightly enriched in mineral-containing conditions both in the presence and absence of tumor cells **(Fig. 4B)**. Since dendritic cells are essential innate immune system responders to biomaterial implants^59^, this increase may be caused by varied host responses at the stiffer M-Bone scaffold interface. Alternatively, the HA content of scaffolds can directly alter or enhance dendritic cell maturation^48^. As some tumor-immune microenvironments can direct dendritic cell differentiation into osteoclasts that drive metastasis-associated bone resorption^60^, our results may indicate that bone matrix changes could affect metastatic progression via differential dendritic cell recruitment or phenotypic changes.

To characterize the effect of mineral on macrophage phenotype in more detail, we first compared macrophages in tumor-free M-Bone and DM-Bone explants. Consistent with epidemiological data and our findings that decreased bone matrix mineral content generates a microenvironment that is more permissive to secondary tumor formation, macrophages on DM-Bone expressed more *Arg1.* The post-translational product of *Arg1*, arginase, is responsible for catabolism of L-arginine into urea and L-ornithine, an amino acid required for cell proliferation and collagen synthesis^61^. Because *Arg1 is* a canonical marker of M2 polarization, which suppresses inflammation and antitumor immunity, our results imply that reduced bone mineral content stimulates macrophage polarization into a phenotype that is conducive to tumor growth^51^ (**Supp.** Fig. 8A). The detected M2-biased macrophages also expressed higher levels of *Fn1,* another indicator of M2 polarization, as well as *Cd163,* and *Mmp9* **(Fig. 4C, 4D)**, which have been associated with worse patient prognosis^62,63^. Differential gene expression and gene ontology analysis of monocyte cells on tumor-free scaffolds confirmed enrichment of genes associated with arginine transport on DM-Bone scaffolds as compared to M-Bone scaffolds (**Supp Fig. 7A, 7B**).

Similar to our findings in the immunocompromised mouse model, absence of mineral enhanced pro-tumorigenic stromal cell traits in tumor-implanted Balb/C mice. More specifically, 4T1.2 tumors growing in mineral-free DM-Bone scaffolds contained more tumor-associated macrophages (TAMs) than tumors in mineral-containing M-Bone scaffolds. These TAMs were characterized by co-expression of the canonical M1 macrophage marker, *Nos2*, and markers of inflammation *Ccl5* and *Il1b* **(Fig. 4E, 4F).** These TAM cells also upregulated the M2 marker *Arg1* as well as the matrix remodeling genes MMP12 and MMP13 and cell adhesion and migration molecule CXCL3. Although M1 phenotypes are thought to be tumor-suppressing, and elevated expression of *Arg1* in cells resembling M1 macrophages seems counterintuitive, macrophages can co-express both markers when degrading and endocytosing collagen I^64^, the primary ECM component in our scaffolds. Indeed, tumors growing in DM-Bone scaffolds were enriched in macrophage phenotypes driving both inflammation and matrix remodeling, processes known to stimulate tumor outgrowth and invasion **(Fig. 4B)**. This mixed macrophage polarization phenotype has been observed in solid tumors *in vivo,* including breast cancer^65,66^, and has been associated with worse outcomes^67^. Together these data indicate that reduced bone mineral content may bias macrophages into an M2-like, anti-inflammatory phenotype when no tumor cells are present. Lack of bone mineral in the presence of tumor cells alters macrophage activation and polarization state further and biases macrophages to be even more pro-tumorigenic.

## DISCUSSION

Decreased bone mineral content is a risk factor for bone metastasis and associated with stromal changes^7,68,69^, but the functional link between bone mineral content and stromal heterogeneity in the presence and absence of tumor cells remains unexplored. Our results suggest that, even in the absence of tumor cells, bone matrix demineralization may prime stromal cells to phenotypes that are supportive of tumor growth regardless of mouse model. The respective mouse model, however, will influence whether fibroblasts or immune cells are the primary mediators of these effects. These findings demonstrate the importance of careful consideration of matrix, mouse, and tumor models when conducting bone metastasis studies, and may help explain why low bone mineral density increases the risk of bone metastasis.

Although ECM modifications and stromal cell phenotype are contributing factors to breast cancer progression^47,70–72^, their importance in bone metastasis is not well characterized in part due to a lack of model systems offering precise control over bone ECM properties. As many conditions comorbid to breast cancer, such as aging, chemotherapy, decreased exercise, hormone changes, and radiation lead to bone mineral decline^4–6^, understanding these connections is critical. While bone mineral density decline can be induced in mice, these models are accompanied by broad systemic effects that impact stromal cell behavior independent of bone matrix^19–21^. Here, we used implantable scaffolds that capture the structure and composition of bone while offering selective control of mineral levels *in vivo*. Leveraging these scaffolds in conjunction with two well accepted and widely used models of triple-negative breast cancer and single-cell RNA-sequencing, we determined that the stromal and immune microenvironment vary significantly in response to changes in bone mineral density, with implications for tumor growth.

Bone-resident stem and stromal cells are known regulators of the outgrowth and aggressiveness of disseminated breast cancer cells, and in other metastatic sites, fibroblasts play critical roles as both tumor-suppressors and tumor-promoters^34,36^. For example, mineralizing culture substrates with HA can affect the differentiation and phenotype of mesenchymal stem cells (MSCs) and their progeny^73,74^, but little is known about how these effects may impact tumor growth.

Previous work with polymeric and urinary bladder ECM implants following muscle injury^23^ revealed new biomaterial signaling modules and cell subsets not previously implicated in response to biomaterials. Expanding upon these findings, our results indicate that bone mineral further regulates host responses to implanted biomaterials and that these changes regulate tumor growth. More specifically, our data suggest that bone matrix devoid of mineral can induce a profibrotic myofibroblastic phenotype, which worsens in the presence of tumor cells. Consistent with clinical evidence that myofibroblasts impair cancer outcomes^75^, this polarization correlated with more rapid tumor growth. Moreover, the fibrotic response to biomaterial implants is correlated with changes in myeloid cell phenotype^76^. Therefore, our findings could have clinical implications as bone matrix-derived, demineralized implants are often used to repair skeletal defects^77^. Our transcriptomic data was corroborated by the increased detection of alpha-SMA+cells by IHC as well as increased collagen deposition characteristic of myofibroblast activity. Although *Acta2*+ CAFs were the dominant phenotype in the absence of an active immune system, we captured few fibroblasts in our immunocompetent model. This could be due to the rapid growth of the 4T1.2 cells, where the driving microenvironmental factor for these tumors is likely necrosis or hypoxia rather than matrix and stromal cell interactions. Thus, while this model enabled examination of immune cells, a limitation of the immunocompromised model, it lacks fibroblast involvement which is a common component of human tumors and bone metastases^34,55,78^. Accordingly, future studies will need to look into these results in other models and explore mechanisms that control increased fibroblast recruitment in bone.

Immune cell state, particularly macrophage polarization state, is a similarly important regulator of tumor growth at metastatic sites^79^, and, according to our results, is influenced by bone mineral content. Prior work indicates that macrophage activation and subsequent polarization to M1 or M2-biased phenotypes, a simplification of the broad spectrum of macrophage transitional states, dictates whether microenvironments are pro- or anti-tumorigenic^79^. For example, cell-cell crosstalk, tumor cell-secreted cytokines, and ECM mechanical properties and sequestered factors have all been found to alter macrophage polarization^80–82^. We now show that in the absence of tumor cells, bone matrix devoid of bone mineral directs macrophages towards M2-biased states which have been associated with worse patient outcomes^83,84^. In mineral-free implants with tumor cells, however, we observed transitional-activated macrophages that co-expressed both M1 and M2 markers and genes regulating matrix remodeling and adhesion. Notably, while *Arg1* is a canonical marker of M1 polarization^84^, its post-translational product, arginase, is also involved in regulating biosynthetic pathways controlling the metabolism of alternatively activated macrophages^85^ in a process that depends on collagen degradation and subsequent endocytosis^64^. Since absence of mineral exposes more of the underlying collagen matrix to cells, it is possible that the *Nos2*+ cells we observed are a profibrotic TAM subtype characterized by high nitric oxide levels induced by increased intracellular arginine from collagen catabolism^86^. Implantation of biomaterial scaffolds into Balb/C mice has been shown previously to recruit immune cells^53,76,87^, and that altering biomaterial properties affects immune cell recruitment. Our results are consistent with these findings **(Fig. 3F)**, and warrant consideration of matrix composition for future site-specific studies of immune cells in metastasis.

In this study we used triple-negative mammary cancer cells, although bone metastases are often found in patients with estrogen receptor-positive (ER^+^) breast cancers. We chose these cells as they are widely used to study bone metastasis in mice^43^, and we have validated that ER^+^ cells are similarly responsive to mineral changes^2^. Nevertheless, recurrence in the bone is more common in patients with ER^+^ breast cancer^88^. Our findings suggest that changes in bone mineral density could contribute to these differences, as triple-negative breast cancer occurs more frequently in younger women with physiological bone mineral density while ER^+^ breast cancer develops more frequently in post-menopausal women, whose reduced estrogen levels decrease bone mineral density^89^. Given these connections, follow-up studies are necessary to compare the role of bone mineral density on the cellular composition of metastatic environments and thus, metastasis formation in different cancer subtypes.

While implanting decellularized bone scaffolds into the flank of mice allowed us to interrogate the effect of bone mineral content on tumor heterogeneity and growth under standardized conditions *in vivo*, several bone-resident cell types could not be considered. However, some of these cell types, such as bone marrow-derived MSCs, can be recruited to implanted biomaterials and tumors via the bloodstream. As HA can drive osteoblast (OB) differentiation, we analyzed our transcriptomic data for OB markers, but did not observe any. This absence could be because MSCs were not recruited to our implants, the short implantation time, or because MSCs differentiated into other cell types, such as CAFs. As both MSC-derived OBs and CAFs play important roles in metastatic lesion progression^30,90^, future studies using bone marrow transplant models to track the fate of bone-marrow derived cells in our implants should study their contribution to tumor growth as a function of matrix mineral content^91^. Moreover, additional work will need to validate the importance of our results with more clinically relevant models of impaired bone mineralization, such as following anti-estrogen therapy^5^, chemotherapy^4^, or vitamin D deficiency^7^. Approaches known to encourage healthy bone formation and mineralization, including mechanical loading^92,93^, could then be tested to see if any such phenotypes can be reversed to prevent tumor growth.

Although the models used here had some limitations, our findings corroborate data collected from clinical bone metastasis samples and thus may offer insights for future investigations. We compared our results to those of an EGFR^-^ (ER^+^/PR^-^/HER2^-^) breast cancer that metastasized to the tibia and pelvis^55^. scRNA-seq of these metastases yielded six cell types, and like in our immunocompromised data, contained predominantly fibroblasts **(Supp.** Fig. 5A). Within the stromal cells, nine fibroblast clusters with diverse functions were identified and a population of *ACTA2*+ *MCAM*+ myofibroblasts as well as a population of myCAF cells were detected across the bone metastases, similar to the *Acta2*+ myCAF-like fibroblasts, which were enriched in the tumor microenvironments on our mineral-free scaffolds **(Supp.** Fig. 5B-D**, 8C)**. Hence, specific fibroblast populations and ACTA2+ CAFs, in particular, affect tumor growth in both our model and clinical bone metastasis samples, underscoring the relevance of our findings to human disease. Future work should aim to evaluate the immune component of bone metastases in a similar manner.

In conclusion, our results suggest that bone matrix changes not only affect the early-stage development and growth of bone lesions via direct interactions with tumor cells, but also that bone demineralization may alter lesion outgrowth via interactions with stromal cells and subsequent changes to the microenvironment **(Fig. 5)**. The finding that lack of bone mineral alone can induce a profibrotic state in stromal and immune cells motivates additional research and early clinical intervention to prevent a decrease of bone mineral density resulting from primary cancer treatments like radiation, hormone therapy, and chemotherapy. Additionally, this finding corroborates that efforts to maintain or increase bone mineral density, such as mechanical loading of bones, may not only be a strategy to interfere with bone metastasis formation in breast cancer patients, but may also improve bone engraftment and defect repair in regenerative settings^93,94^. Collectively, our results motivate careful consideration of the heterogeneous and multicellular responses to bone matrix changes in metastasis and disease and could yield new avenues of research toward clinical intervention that could improve bone defect repair or bone disease treatment and slow bone metastatic progression.

**Figure 5.**
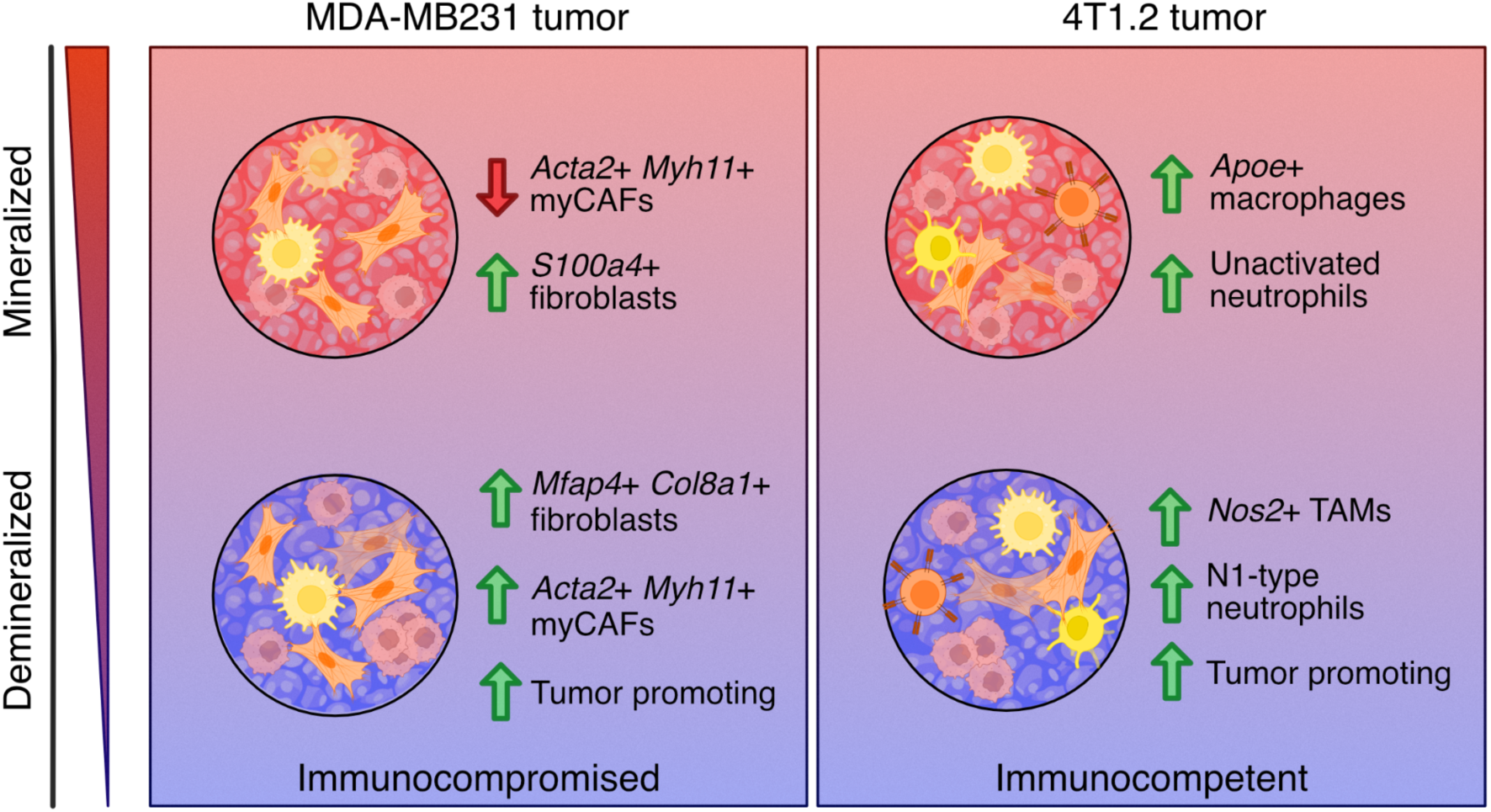
Reduced bone matrix mineral content induces global changes to the tumor microenvironment. Microenvironmental changes caused by reduced mineral density include 1) the induction of proinflammatory, myofibroblastic, and pro-tumorigenic conditions via the induction of myofibroblastic cancer-associated fibroblasts (myCAFs) in the absence of immune activation, and 2) the induction of complex tumor-promoting and pro-inflammatory conditions, induced by tumor-associated macrophages (TAMs) with a mixed-polarization phenotype and activated tumor-associated neutrophils (TANs) in the presence of a competent immune response.

## METHODS

### Fabrication of bone matrix scaffolds

M-Bone and DM-Bone bone matrix scaffolds were fabricated as previously described^2^. Briefly, 6 mm diameter cylindrical subchondral bone plug biopsies were extracted from 1-3 day old neonatal bovine femurs, flushed with deionized water to remove marrow, and cut into 1 mm thick sections. Cells and debris were removed by incubation in an extraction buffer of 20 mM NaOH and 0.5% Triton X-100 in PBS at 37 °C. Scaffolds were then treated with 20 U/mL DNase I to remove residual DNA, washed 5 times with PBS and labeled as “M-Bone’’. To produce demineralized “DM-Bone” scaffolds, mineral-containing “M-Bone” scaffolds were demineralized at physiological pH in a solution of 9.5% ethylenediaminetetraacetic acid (EDTA), then washed 5 times in PBS. Before use, scaffolds were sterilized by overnight incubation in 70% ethanol, then washed with PBS and pre-incubated in complete culture medium.

### Materials characterization of bone matrix scaffolds

For characterization of bone matrix scaffolds, M-Bone and DM-Bone samples were prepared and excess moisture was removed by aspiration. Mechanical testing of scaffolds was performed by dynamic mechanical thermal analysis (DMA Q800, TA Instruments, US) in compression mode. Samples were loaded between the two plattens of the clamp with a preload force of 0.1 N. M-Bone and DM-Bone scaffolds were compressed at ambient temperature and pressure with force ramps of 0.1 N/min and 2 N/min to a threshold of 0.8 N and 18 N, respectively. Scaffold mineralization and structure was assessed by micro-computed tomography using a Skyscan 1276 (Bruker, DE). Scaffolds were fixed with 4% PFA and contrast enhanced with diffusible-iodine in a 0.5% solution in buffered Lugol solution, then scanned with a voltage of 100 kV, using an Al and Cu filter, at a resolution of 10 microns per pixel, reconstructed using the standard Bruker reconstruction software, and false-colored on the basis of attenuation coefficient (Avizo, ThermoFisher, US). Scaffold trabecular structure and collagen fiber structure were assessed by scanning electron microscopy (Gemini 500, Zeiss, DE). Samples were fixed with 2% glutaraldehyde in 0.05M sodium cacodylate buffer, stained for 1 hour with 1% (w/v%) OsO_4_, and sequentially dehydrated by ethanol series before CO_2_ critical point drying for 48 hours. Samples were carbon coated (Desk II, Denton Vacuum, US) and imaged in secondary electron mode (SE) with an electron voltage of 10 kV.

### Cell culture and implant preparation

MDA-MB231 breast cancer cells (ATCC) expressing RFP and luciferase were routinely cultured in DMEM (ThermoFisher Scientific, US) supplemented with 10% fetal bovine serum (FBS) (Atlanta Biologicals, US) and 1% penicillin/streptomycin (P/S) (ThermoFisher Scientific, US) in a 5% CO2 incubator. 400,000 cells (P12) were seeded onto M-Bone and DM-Bone scaffolds and cultured overnight before implantation. Bone-tropic Balb/C-syngeneic 4T1.2 cells expressing GFP and luciferase were routinely cultured in RPMI-1640 (ThermoFisher Scientific, US) supplemented with 10% fetal bovine serum (FBS) (Atlanta Biologicals, US) and 1% penicillin/streptomycin (P/S) (ThermoFisher Scientific, US) in a 5% CO2 incubator. 50,000 cells (P4) were seeded onto M-Bone and DM-Bone scaffolds and cultured overnight before implantation.

### Animal experiments

Mouse experiments were conducted following Cornell University animal care guidelines and were approved by Cornell University’s Institutional Animal Care and Use Committee. All animals received 4 scaffolds implanted on the subcutaneous flank: 1 unseeded DM-Bone, 1 unseeded M-Bone, 1 pre-seeded DM-Bone, and 1 pre-seeded M-Bone scaffold. For immunocompromised experiments, scaffolds were implanted on the subcutaneous flank of 6–8-week-old female athymic nude-Foxn1^nu^ mice (Envigo, US). Bioluminescence images were taken with an *in vivo* imaging system (IVIS Spectrum, Perkin Elmer, US) 10 minutes after intraperitoneal (IP) injection of 150 mg/kg D-luciferin (Gold Biotechnology, US) in DPBS once a week. For immunocompetent experiments, pre-seeded scaffolds were implanted on the subcutaneous flank of 6–8-week-old female Balb/C mice. Bioluminescence images were taken 15 minutes following IP injection of 150 mg/kg D-luciferin in DPBS twice a week. Implants were followed for 14 days and candidates for sequencing were identified by bioluminescent signal. Samples for single-cell transcriptomics were processed as below. Other samples were fixed in 4% paraformaldehyde (PFA) overnight at 4 ℃, washed with PBS, stored in 70% EtOH, and paraffin-embedded.

### Histology and staining

Explanted scaffolds were fixed with ice-cold 4% paraformaldehyde and decalcified with 10% EDTA. Paraffin sections were used for H&E staining as well as Masson’s Trichrome staining and immunohistochemical staining of αSMA and F4/80. For IHC stains, sections were deparaffinized, and subjected to antigen retrieval in citrate buffer at 95℃ for 20 minutes. After extraction, samples were blocked with horse serum, then incubated with either primary rabbit anti-mouse αSMA antibody (ab124964, Abcam) or primary rat anti-mouse F4/80 antibody (ThermoFisher, clone 14-4801-82). Primaries were detected with biotinylated horse anti-rabbit and rabbit anti-rat secondaries (Vectorlabs), respectively, then incubated with a streptavidin-peroxidase tertiary antibody and developed with stable peroxidase substrate solution (ThermoFisher). All sections were counterstained with Mayer’s or Harris hematoxylin (ThermoFisher) and imaged using a Scanscope slide scanner (Aperio CS2, Leica Biosystems) with a 40x objective. To quantify the collagen content of Masson’s Trichrome stained sections, images of trichrome stained sections were uploaded to QuPath v.0.2.0^95^ and split into 3 vector channels (0.762, 0.609, 0.222) then segmented into a grid. Trabeculae and white space were excluded, pixel intensity in the collagen channel was calculated for each ROI.

### Sample preparation for single-cell transcriptomics (MDA-MB-231)

Mice implanted with scaffolds were sacrificed on day 14 post implantation and the scaffolds with tumor microenvironment were isolated aseptically for single-cell RNA-sequencing experiments. One half of the collected scaffolds from each condition was minced into 1-2 mm pieces and transferred to 1.5 ml Eppendorf tube for dissociation into a single cell suspension. Tissue pieces were then digested in a tissue dissociation media with 5 mg/mL dispase, 2.5 mg/mL collagenase I, and 2.5 mg/mL collagenase II in 1x basal culture medium in a 37°C incubator. At the end of the digestion, the cells were passed through a 40 µm filter and centrifuged into a pellet. To remove erythrocytes from the suspension, samples were resuspended in an ammonium-chloride-potassium (ACK) lysis buffer (Lonza #10-548E) for 3-5 minutes and centrifuged at 180 g for 6 minutes. Samples were then washed 3x in PBS with 0.04% BSA and then resuspended at 1 × 10^6^ cells per ml. Cells from each sample were stained with Trypan Blue and cell viability was calculated on an automated cell counter (Countess II) before loading the cells on 10x Chromium. We used these cell viabilities to adjust the number of cells loaded on 10x Chromium to get the desired number of transcriptomes from viable cells for each sample (5000 cells per sample).

### Sample processing for single-cell RNA-sequencing with Cell Multiplexing (4T1.2)

Mice implanted with scaffolds were sacrificed on day 14 post implantation and the scaffolds with tumor microenvironment were isolated aseptically for single-cell RNA-sequencing experiments. One half of the collected scaffolds from each condition was minced into 1-2 mm pieces and transferred to a 1.5 ml Eppendorf tube for dissociation into a single cell suspension. Tissue pieces were then digested in a tissue dissociation media with 5mg/mL dispase, 2.5 mg/mL collagenase I, and 2.5 mg/mL collagenase II in 1x basal culture medium in a 37°C incubator. At the end of the digestion, the cells were passed through a 40 µm filter and centrifuged into a pellet. To remove erythrocytes from the suspension, samples were resuspended in an ammonium-chloride-potassium (ACK) lysis buffer (Lonza #10-548E) for 3-5 minutes and centrifuged at 180 g for 6 minutes. Samples were then washed twice in PBS with 0.04% BSA and then resuspended at 0.5 × 10^6^ cells in 1 ml total buffer. The cells were then centrifuged, and the supernatant was removed without disturbing the pellet. Distinct 10x Genomics Cell Multiplexing Oligos were then added to all samples and incubated for 5 minutes at room temperature. Samples Control-M-Bone, Control-DM-Bone, Tumor-M-Bone, and Tumor-DM-Bone were tagged with CMO301, CMO302, CMO303, and CMO304 respectively. Labeled cell suspensions were then washed 3x in pre-chilled wash and resuspension buffer containing 1% BSA in 1x PBS as recommended in the manufacturer’s Cell Multiplexing Oligo Labeling protocol for Single-cell RNA Sequencing. The cells were resuspended in the resuspension buffer at 1500 cells/ul and were counted on an automated cell counter (Countess II). Labeled cells from individual samples were pooled in equal proportions, stained with Trypan Blue, and cell viability was calculated on an automated cell counter (Countess II) before loading the cells on 10x Chromium. We used these cell viabilities to adjust the number of cells loaded on 10x Chromium to get the desired number of transcriptomes from viable cells for each sample (4,000 cells per sample; 16,000 total cells for four samples).

### Single-cell RNA-sequencing library preparation (MDA-MB-231)

Single-cell suspensions were loaded on the Chromium platform (10x Genomics) with one microfluidic channel per condition and ∼5000 target cells per channel. Single-cell mRNA libraries were made using the Chromium Next GEM Single Cell 3’ Library Construction V3.1 Kit according to the manufacturer’s protocol. Sequencing Libraries sequenced on an Illumina NextSeq 500/550 using 75 cycle high output kits (Index 1 = 8bp, Index2 =8bp, Read 1 = 28, and Read 2 = 55bp). Raw sequencing data was aligned to a combined human and mouse genome reference (assembly: GRCh38 and mm10) using the Cell Ranger 6.1.1 pipeline (10x Genomics) to obtain single-cell gene expression matrices for individual samples.

### Single-cell RNA-sequencing library preparation (4T1.2)

Oligo-tagged and pooled single-cell suspension derived from all four experiment conditions was loaded on a single microfluidic channel of the Chromium platform (10x Genomics) with a target of ∼16,000 cells (∼4,000 cells per experiment condition). A single-cell mRNA library and a cell multiplexing oligo library were made using the Chromium Next GEM Single Cell 3’ Library Construction V3.1 Kit with Feature Barcode technology for Cell Multiplexing, according to the manufacturer’s protocol. Sequencing Libraries sequenced on an Illumina NextSeq 500/550 using 75 cycle high output kits (Index 1 = 10 bp, Index2 =10 bp, Read 1 = 28 bp, and Read 2 = 90 bp). Raw sequencing data was aligned to a combined human and mouse genome reference (assembly: GRCh38 and mm10) using the Cell Ranger 6.1.1 pipeline (10x Genomics).

### scRNA-seq data preprocessing, analysis, and visualization (MDA-MB-231)

Raw gene expression matrices for individual samples were loaded in an R-4.0.3 environment using Seurat package (v4.3.0)^96^. After excluding the cells with less than 200 unique genes, less than a 1,000 unique transcript UMIs, or more than 30 percent mitochondrial transcripts, we analyzed 5063, 4822, 3401, and 3687 single-cell transcriptomes from Control-M-Bone, Control-DM-Bone, Tumor-M-Bone, and Tumor-DM-Bone, respectively. The 16,973 cells across four experiment conditions were then normalized and log-transformed using the Seurat package. We used the FindVariableFeatures function in Seurat to choose the top 2000 highly variable genes from the dataset using the “vst” selection method. We then performed mean centering and scaling, followed by principal component analysis (PCA) to reduce the dimensions of the data to the top 20 principal components. t-Distributed Stochastic Neighbor Embedding (t-SNE) was initialized in this PCA space to visualize the data on reduced t-SNE dimensions. The cells were clustered on PCA space using the Louvain algorithm on a k-nearest neighbors graph constructed using gene expression data as implemented in FindNeighbors and FindClusters commands in the Seurat package. Cell-type-specific canonical gene markers were used to label cell clusters differentially expressing those markers. To accurately label individual clusters, differentially expressed genes for each cluster were found with the FindAllMarkers command using the Wilcox test. Single-cell gene expression was visualized using FeaturePlot, DoHeatMap, and VlnPlot functions from Seurat.

### scRNA-seq data preprocessing, analysis, and visualization (4T1.2)

Cell multiplexing oligo sequencing data was used to demultiplex the raw gene expression data for individual samples using the multi command in Cell Ranger 6.1.1 pipeline. The cell multiplexing oligo IDs were provided in a config file to the CellRanger multi command and min-assignment-confidence was set to 0.9 (recommended default). Raw gene expression matrices for cellplex samples were loaded in an R-4.0.3 environment using Seurat package (v4.3.0). After excluding the cells with less than 200 unique genes, less than a 1,000 unique transcript UMIs, or more than 30 percent mitochondrial transcripts, we analyzed 2082, 1649, 2270, and 3252 single-cell transcriptomes from Control-M-Bone, Control-DM-Bone, Tumor-M-Bone, and Tumor-DM-Bone, respectively. Single-cell transcriptomes were preprocessed, analyzed, and visualized similar to the analysis for MDA-MB-231 cells discussed above.

### scRNA-seq data preprocessing, analysis, and visualization for human bone metastases

Single-cell gene expression count matrices and meta data files were downloaded from Gene Expression Omnibus and loaded directly in an R-4.0.3 environment. The single cell transcriptomes from 10,056 cells across two human bone metastases samples were then processed using Seurat package (v4.3.0). We used the FindVariableFeatures function in Seurat to choose the top 2000 highly variable genes from the dataset using the “vst” selection method. We then performed mean centering and scaling, followed by principal component analysis (PCA) to reduce the dimensions of the data to the top 20 principal components. t-Distributed Stochastic Neighbor Embedding (t-SNE) was initialized in this PCA space to visualize the data on reduced t-SNE dimensions. Cell type labels provided in the meta data were directly used to identify all cell types and to isolate stromal fibroblast cells. Single-cell gene expression for genes of interest was visualized using FeaturePlot functions in the Seurat package.

## DATA AVAILABILITY

The authors declare that all sequencing data supporting the findings of this study have been deposited in NCBI’s Gene Expression Omnibus (GEO) with GEO series accession number <u>GSE256109</u>. scRNA-seq datasets from metastatic human breast cancer were downloaded from GEO repository: <u>GSE190772</u>. All other data supporting the findings in this study are included in the main article and associated files.

## CODE AVAILABILITY

Scripts to reproduce the analysis presented in this study have been deposited on GitHub (https://github.com/madhavmantri/bone_matrix).

## ACKNOWLEDGEMENTS

We would like to thank Dr. Peter Schweitzer and the Cornell Genomics Center (RRID:SCR_021727) for help with sequencing assays and the Cornell Bioinformatics facility for assistance with bioinformatics. We also thank the Cornell Animal Health Diagnostic Core for paraffin embedding and sectioning and the Cornell Center for Animal Resources and Education (CARE) staff for animal care. We thank Dr. Lawrence Bonassar for providing materials to produce the bone scaffolds. We thank the members of the Fischbach and De Vlaminck labs for many valuable discussions. This work was supported by a seed grant provided to C.F and I.D.V. by the NCI through the Center on the Physics of Cancer Metabolism (1U54CA210184) and NSF GRFP (DGE-1650441) to M.A.W. This work used the Cornell Center for Materials Research, which is supported through the NSF MRSEC program (DMR-1719875), and the Cornell University Biotechnology Resource Center (BRC) facilities, including the IVIS Spectrum (NIH S10OD025049) and the SkyScan 1276 mouse CT (NIH S10OD025049).

## AUTHOR CONTRIBUTIONS

M.W., M.M., C.F., and I.D.V. designed the study. M.W. and M.M. performed the animal experiments. M.W., M.M., and E.S. performed the single-cell transcriptomics experiments and analyzed the scRNA-seq data. M.W. performed histology and analyzed the images. M.W., M.M., I.D.V., and C.F wrote the manuscript. All authors provided feedback and comments.

## MATERIALS & CORRESPONDENCE

Please direct correspondence to Claudia Fischbach and Iwijn De Vlaminck.

