## Supplementary Information for "Bone mineral density affects tumor growth by shaping microenvironmental heterogeneity"

**AFFILIATIONS**:
[1]Nancy E. and Peter C. Meinig School of Biomedical Engineering, Cornell University, Ithaca, New York 14850
[2]Department of Materials Science and Engineering, Cornell University, Ithaca, NY 14850
[3]Kavli Institute at Cornell for Nanoscale Science, Cornell University, Ithaca, NY 14850

*These authors contributed equally

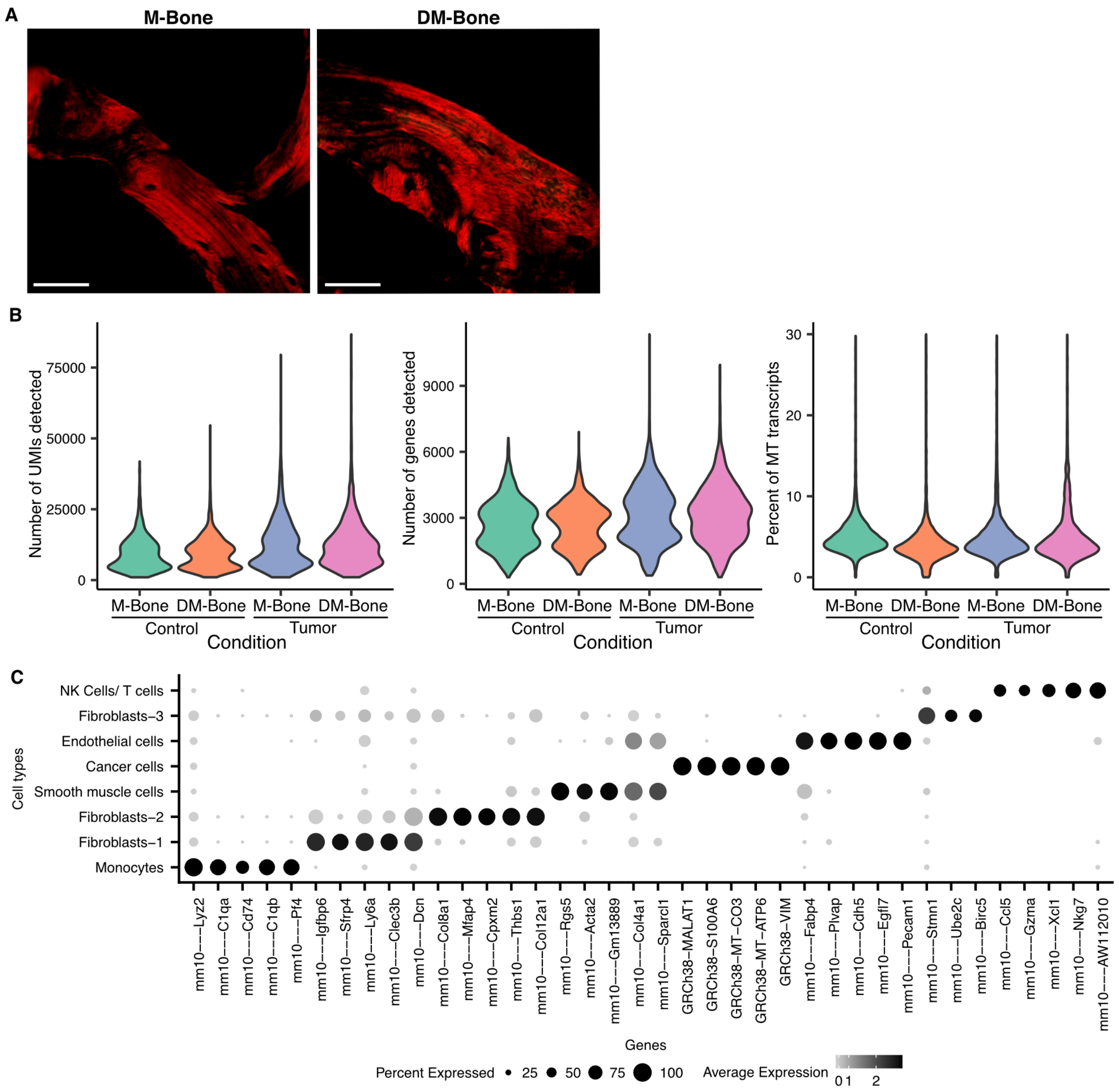


**Supplementary Figure 1: Single-cell transcriptomics of tumor-free and MDA-MB231 tumor cell-containing M-Bone and DM-Bone scaffolds. A)** Representative polarized-light images of Picrosirius Red stained sections of M-Bone and DM-Bone scaffolds. Scale bar = 50 µm **B)** Number of unique transcripts per cell (left), number of unique genes detected per cell (center), and percentage of mitochondrial transcripts (right) in scRNA-seq datasets across tumor-free and tumor-containing M-Bone and DM-bone scaffolds. **C)** Top-five differentially expressed genes (two-sided Wilcoxon test, log2 fold-change > 1.0 and p-value < 0.01) for all cell types across four experiment conditions.

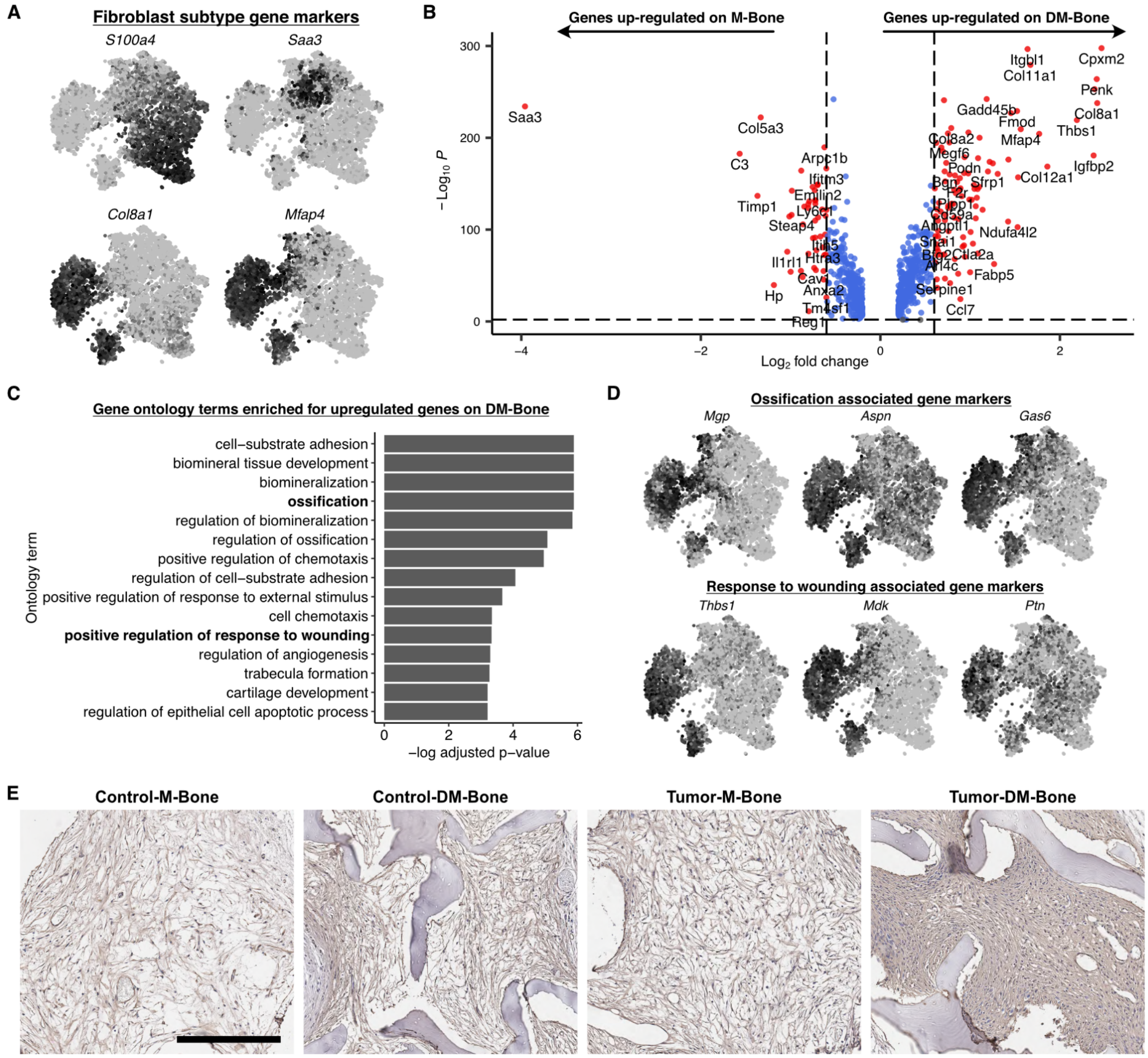


**Supplementary Figure 2: Stromal fibroblast phenotype is altered by bone matrix mineral content. A)** t-Distributed Stochastic Neighbor Embedding (t-SNE) feature plots showing expression of gene markers for fibroblast subtypes detected in scRNA-seq datasets on tumor-free and tumor-containing M-Bone and DM-Bone scaffolds. **B)** Volcano plot showing differential gene expression analysis results on comparison of fibroblast cells on tumor-free M-Bone and DM-Bone scaffolds. **C)** Bar plot showing top 15 gene ontology terms enriched in genes upregulated in fibroblast cells on tumor-free DM-Bone scaffolds (Control-DM-Bone) as compared to M-Bone scaffolds. (Control-M-Bone). **D)** t-SNE feature plots showing expression of three genes associated with ossification: *Mgp*, *Aspn*, and *Gas6*; and three genes associated with response to wound healing: *Thbs1*, *Mdk1*, *Ptn*. **E)** Representative images of anti-α-Smooth muscle actin immunohistochemical staining of explanted scaffolds. Scale bar = 300 µm.

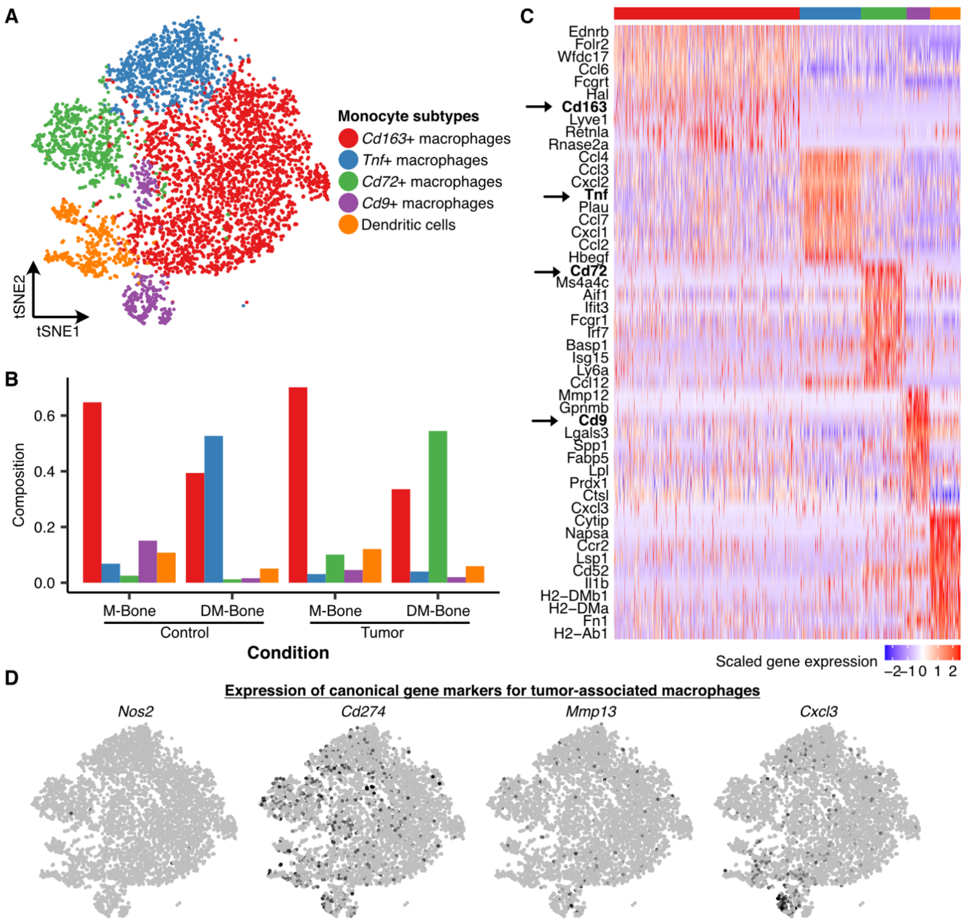


**Supplementary Figure 3: Myeloid cell heterogeneity in MDA-MB231 microenvironment on M-Bone and DM-Bone scaffolds. A)** t-Distributed Stochastic Neighbor Embedding (t-SNE) map of 5,866 monocyte single-cell transcriptomes clustered by gene expression and colored by the monocyte subtype labels. **B)** Bar plot showing relative proportion of various monocyte cell clusters across the four experimental conditions. Colors represent the fibroblast subtypes as shown in Supp. Fig. 3A. **C)** Heat map showing the log-normalized and scaled expression of top-ten differentially expressed genes in each monocyte cluster. Colors in the color bar on top represent the fibroblast subtypes as shown in Supp. Fig. 3A. **D)** t-SNE feature plots showing expression of markers of tumor-associated macrophages.

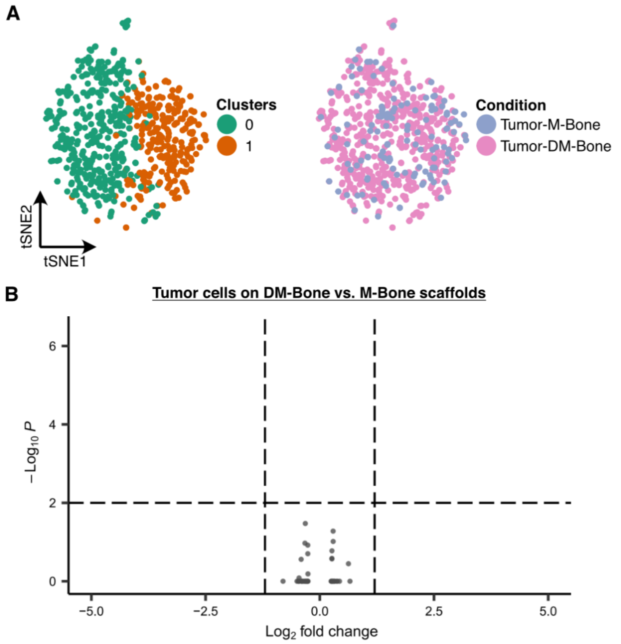


**Supplementary Figure 4: MDA-MB231 heterogeneity between conditions is minimal. A)** t-Distributed Stochastic Neighbor Embedding (t-SNE) map of 649 single-cell transcriptomes of MDA-MB231 cells clustered by gene expression and colored by the cluster ID (left) and scaffold condition (right). **B)** Volcano plot showing differential gene expression analysis results on comparison of MDA-MB-231 tumor cells on M-Bone and DM-Bone scaffolds.

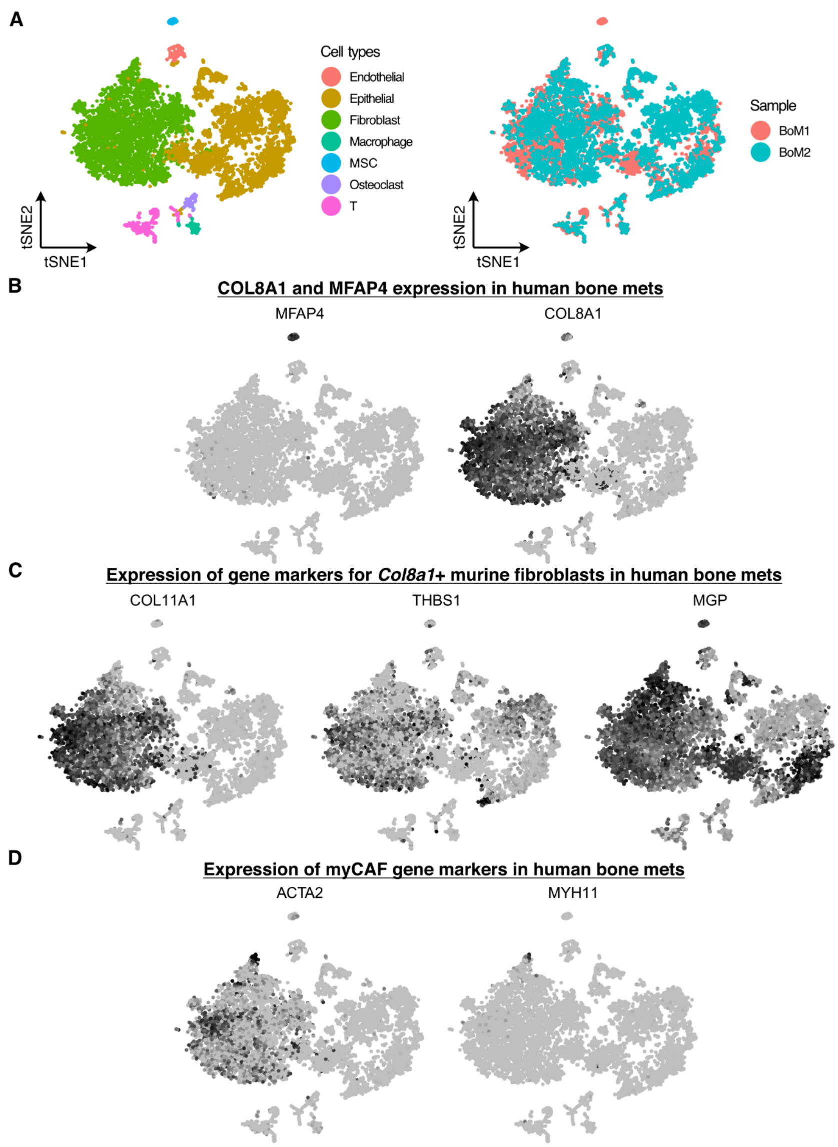


**Supplementary Figure 5: Microenvironment heterogeneity of human bone metastatic tumors.** **A)** t-Distributed Stochastic Neighbor Embedding (t-SNE) map of 10,056 single-cell transcriptomes clustered by gene expression and colored by the cell type (left) and sample ID (right). **B)** t-SNE feature plots showing expression of COL8A1 and MFAP4 in fibroblast cells from human bone metastases. **C)** t-SNE feature plots showing expression of markers of *Col8a1*+ *Mfap4*+ murine fibroblast cells in human bone metastases. **D)** t-SNE feature plots showing expression of markers of myCAFs in human bone metastases.

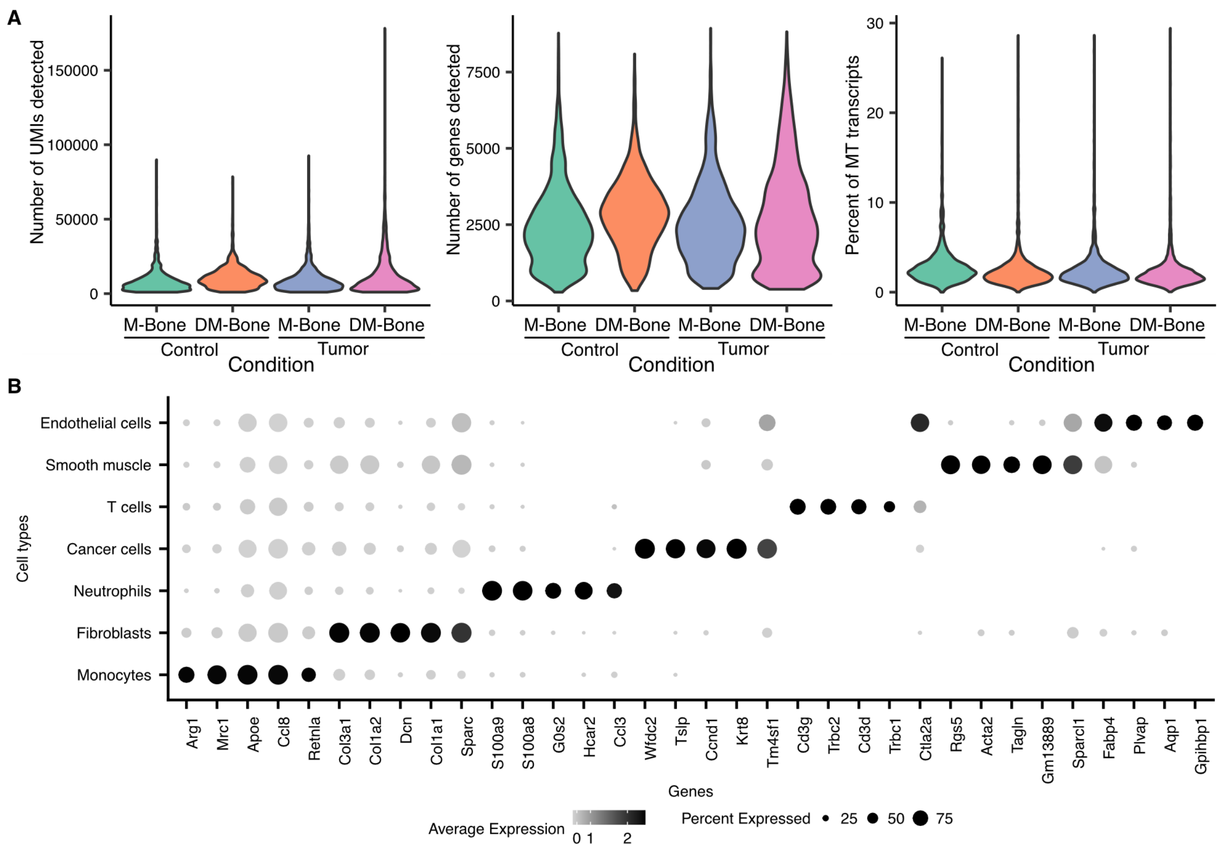


**Supplementary Figure 6: Single-cell transcriptomics of 4T1.2 tumor cell-containing M-Bone and DM-Bone scaffolds. A)** Number of unique transcripts per cell (left), number of unique genes detected per cell (center), and percentage of mitochondrial transcripts (right) in scRNA-seq datasets across tumor-free and tumor-containing M-Bone and DM-bone scaffolds. **B)** Top-five differentially expressed genes (two-sided Wilcoxon test, log2 fold-change > 1.0 and p-value < 0.01) for all cell types across four experiment conditions.

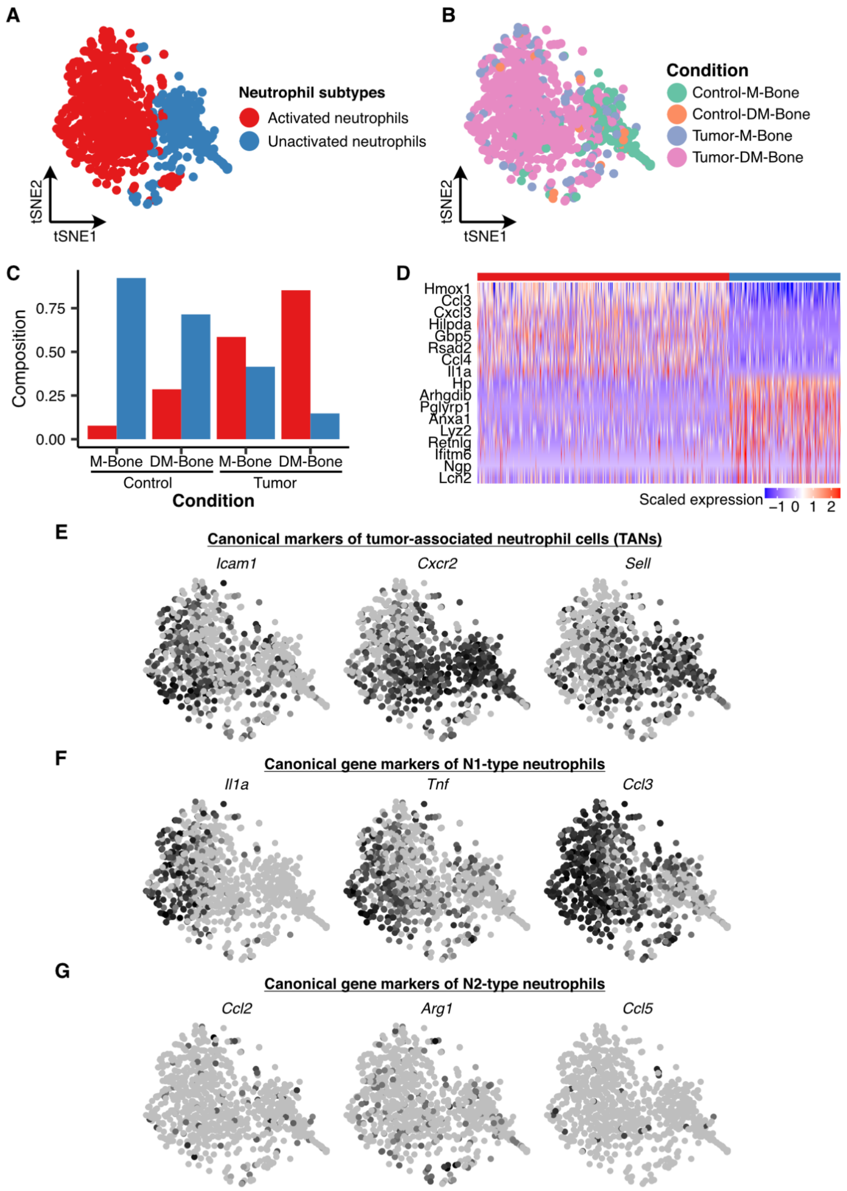


**Supplementary Figure 7: Neutrophil heterogeneity in 4T1.2 microenvironment on M-Bone and DM-Bone scaffolds. A)** t-Distributed Stochastic Neighbor Embedding (t-SNE) map of 754 neutrophil single-cell transcriptomes clustered by gene expression and colored by the neutrophil subtype clusters. **B)** t-SNE map of 754 neutrophil single-cell transcriptomes clustered by gene expression and colored by the experimental conditions. **C)** Bar plot showing relative proportion of various neutrophil cell clusters across the four experimental conditions. Colors represent the neutrophil subtypes as shown in Supp. Fig. 7A. **D)** Heat map showing the log-normalized and scaled expression of top-ten differentially expressed genes in each neutrophil cluster. **E)** t-SNE feature plots showing expression of canonical markers of tumor-associated neutrophils. **F)** t-SNE feature plots showing expression of markers of N1-type neutrophils. **G)** t-SNE feature plots showing expression of markers of N2-type neutrophils.

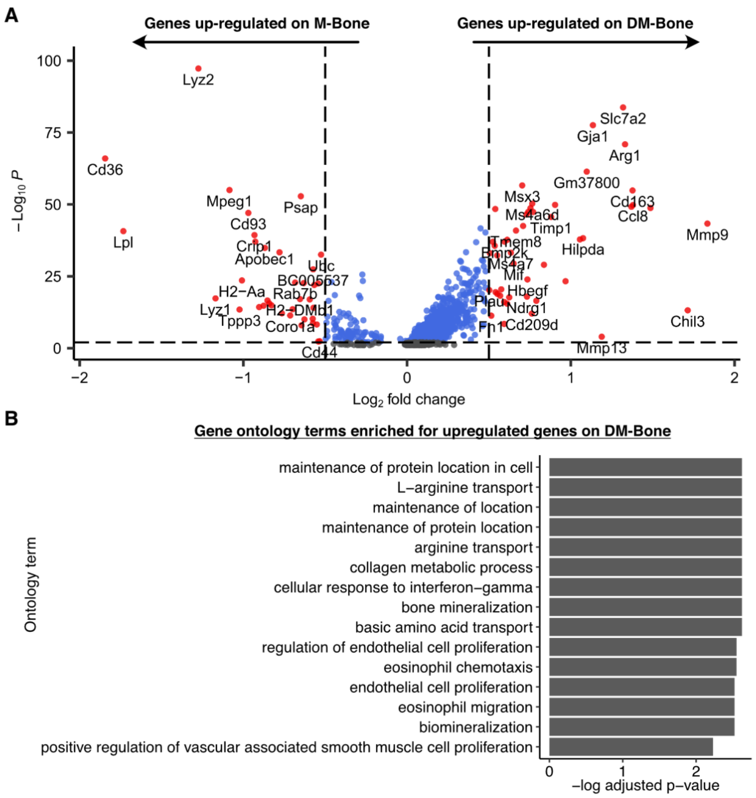


**Supplementary Figure 8: Macrophage activation state is altered by bone mineralization. A)** Volcano plot showing differential gene expression analysis results on comparison of macrophage cells on tumor-free M-Bone and DM-Bone scaffolds. **B)** Bar plot showing top 15 gene ontology terms enriched in genes upregulated in monocyte cells on tumor-free DM-Bone scaffolds as compared to M-Bone scaffolds.
